# The mitochondrial genomes of the mesozoans *Intoshia linei, Dicyema* sp., and *Dicyema japonicum*

**DOI:** 10.1101/282285

**Authors:** Helen. E. Robertson, Philipp. H. Schiffer, Maximilian. J. Telford

## Abstract

The Dicyemida and Orthonectida are two groups of tiny, simple, vermiform parasites that have historically been united in a group named the Mesozoa. Both Dicyemida and Orthonectida have just two cell layers and appear to lack any defined tissues. They were initially thought to be evolutionary intermediates between protozoans and metazoans but more recent analyses indicate that they are protostomian metazoans that have undergone secondary simplification from a complex ancestor. Here we describe the first almost complete mitochondrial genome sequence from an orthonectid, *Intoshia linei*, and describe nine and eight mitochondrial protein-coding genes from *Dicyema* sp. and *Dicyema japonicum*, respectively. The 14,247 base pair long *I. linei* sequence has typical metazoan gene content, but is exceptionally AT-rich, and has a divergent gene order compared to other metazoans. The data we present from the Dicyemida provide very limited support for the suggestion that dicyemid mitochondrial genes are found on discrete mini-circles, as opposed to the large circular mitochondrial genomes that are typical across the Metazoa. The *cox1* gene from dicyemid species has a series of conserved in-frame deletions that is unique to this lineage. Using *cox1* genes from across the genus *Dicyema*, we report the first internal phylogeny of this group.

**Key Findings:**

- We report the first almost-complete mitochondrial genome from an orthonectid parasite, *Intoshia linei*, including 12 protein-coding genes; 20 tRNAs and putative sequences for large and small subunit rRNAs. We find that the *I. linei* mitochondrial genome is exceptionally AT-rich and has a novel gene order compared to other published metazoan mitochondrial genomes. These findings are indicative of the rapid rate of evolution that has occurred in the *I. linei* mitochondrial genome.
- We also report nine and eight protein-coding genes, respectively, from the dicyemid species *Dicyema* sp. and *Dicyema japonicum*, and use the *cox1* genes from both species for phylogenetic inference of the internal phylogeny of the dicyemids.
- We find that the *cox1* gene from dicyemids has a series of four conserved in-frame deletions which appear to be unique to this group.

## Introduction

The Mesozoa is an historic name given to two different groups of very small, vermiform and morphologically simple parasitic animals: the Dicyemida, whose members are made up of approximately 40 cells (Furuya & Tsuneki, 2003); and the Orthonectida, which posses just a few hundred cells. As there is uncertainty over whether Dicyemida and Orthonectida form a monophyletic group, the Mesozoa grouping is now generally used informally rather than as a formal taxonomic assignment as a phylum. Both dicyemids and orthonectids are parasites of various marine animals: the dicyemids live in the renal tissue of cephalopods, whilst orthonectids occupy the internal body spaces of a variety of marine invertebrates, including brittle stars, bivalve molluscs, nemerteans and polychaetes.

Adults of both dicyemids and orthonectids have just two cell layers. Although they are multicellular, they lack defined tissues or organs, and there is no evidence for true ectoderm or endoderm specification (Margulis & Chapman, 2009). The adult animals have an external layer of multiciliated cells, which facilitate movement. Internal to this are one or more reproductive cells, but the degree of further cellular differentiation in dicyemids is unclear. A small nervous system, comprising 10-12 nerve cells, and a simple muscular system, composed of four longitudinal and 9-11 circular muscle cells, have been identified in the orthonectid *Intoshia linei* (Slyusarev & Starunov, 2016). Given their simple body organisation, members of the Mesozoa were described in the 19th century – as their name suggests - as being an evolutionarily intermediate between the protozoans and the metazoans. More recent reassessment of mesozoan species indicates that they are in fact bilaterian metazoans that have undergone extreme simplification from a more complex ancestor (Dodson, 1956). *In situ* hybridisations for 16 diverse genes in different life stages of *Dicyema sp*. suggested the presence of multiple different cell types, and provided further support for the idea of a complex ancestor of mesozoan members, followed by extreme simplification of body organisation (Ogino et al., 2011).

Two significant contributions for understanding the evolutionary history of the Mesozoa are the recent publication of the nuclear genome of *Intoshia linei*, which represents one of approximately 20 species of this genus, and a transcriptome of *Dicyema japonicum* (Lu et al., 2017; Mikhailov et al., 2016). The genomic sequence of *I. linei* is 43 Mbp in length and encodes just ∼9000 genes, including those essential for the development and activity of muscular and nervous systems. Neither a phylogenomic analysis based on 500 orthologous groups, nor an analysis of transcriptomic data from *D. japonicum* and a dataset compiled from 29 taxa and >300 gene orthologues, could confidently place the Orthonectida or Dicyemida in a precise position within the Lophotrochozoa. However, strong statistical support was found for a grouping of the Dicyemida with the Orthonectida in the phylum Mesozoa in one of these recent studies (Lu et al., 2017). Most recently, a broad phylogenomic analysis found *Intoshia* to be nested within the annelids (Schiffer et al. under revision elsewhere) (Schiffer et al., 2017). While the position of Dicyema within the Lophotrochozoa could not be unambiguously resolved, it seems clear that Orthonectids and Rhombozoa are not joined in a single taxon. Nevertheless, these species can be regarded as another case of extreme morphological and genomic simplification found in parasites.

One resource that has not been extensively investigated to study the evolution and biology of the mesozoans is their mitochondrial genomes. Interestingly, the limited mitochondrial data analysed from Dicyemida to date suggest a highly unusual mitochondrial gene structure. The three mitochondrial genes that have been sequenced from *D. misakiense* (*cox1, 2* and *3*) appear to be on individual mini-circles of DNA, rather than being part of a single circular genome containing all genes, as is typical of other mitochondrial genomes (Watanabe et al., 1999; Catalano et al., 2015; Awata et al., 2005). Although the vast majority of metazoan mitochondrial genomes are found as a single circular molecule, a few multipartite circular genomes (that is, genomes where mitochondrial genes are found on more than one closed-circle molecule) have been reported across the Bilateria. The majority of these appear to come from parasitic species, and multipartite mitochondrial genomes have been described, for example, from several species of lice(Cameron et al., 2011; Dong et al., 2014; Shao et al., 2005) (Arthropoda) and parasitic nematodes (Hunt et al., 2016; Phillips et al., 2016). The Dicyemida could therefore represent another example of a parasitic organism with a multipartite mitochondrial genome.

*Cox1* genes have been sequenced from a number of Dicyemida species, (*D. koinonum*; *D. acuticephalum*; *D. vincentense*; *D. multimegalum*; *D. coffinense*) but these have not been used in a published molecular systematic study.

In order to provide new mitochondrial gene data from these species for understanding more about their parasitic biology and phylogenetic history, we looked for mitochondrial sequences in publicly available short read sequence data from members of the genus *Dicyema* (*D. japonicum* and *Dicyema sp.*) and Orthonectida (*Intoshia linei*). We compared features of the mitochondrial genome of *I. linei* to other metazoan members to shed additional light on its possible rapid and extreme simplification, and used mitochondrial protein-coding gene data from *Dicyema* sp. and *D. japonicum* to investigate the internal phylogeny of the dicyemids.

## Materials and Methods

### Genome and transcriptome assemblies

We downloaded genomic (*I. linei*: SRR4418796, SR4418797) and transcriptomic (*Dicyema* sp.: SRR827581; *Dicyema japonicum*: DRR057371) data from the NCBI short read archive. Adapter sequences and low quality bases were removed from the sequencing reads using Trimmomatic (Bolger et al., 2014). The *I. linei* genome was re-assembled using the CLC assembly cell (v.5.0 http://www.qiagenbioinformatics.com/) and the CLC assembly cell and Trinity pipeline (Haas et al., 2013) (v.2.3.2) used to assemble the *Dicyema* sp. and *D. japonicum* transcriptomes, with default settings.

### Mitochondrial genome fragment identification and annotation

Mitochondrial gene protein-coding sequences from flatworms were used as tblastn queries to search for mitochondrial fragments in the *I. linei* genome assembly and *Dicyema* sp. and *D. japonicum* Trinity RNA-Seq assemblies, using NCBI translation table 5 ‘invertebrate mitochondrial’. Positively identified sequences were blasted against the NCBI nucleotide database in order to detect possible contaminant sequences from *Dicyema* host species. For each *Dicyema* sp. gene-bearing contig, we found a number of other contigs that had very high sequence similarity to *Octopus* or other cephalopods, indicating a degree of host species contamination in the RNA-Seq data. These sequences were discarded from subsequent analysis.

From initial blast queries, a 14,176 bp mitochondrial contig was identified from the *I. linei* reassembled genome. An additional mitochondrial contig was found that overlapped with this contig to extend the mitochondrial sequence of *I. linei* to 14,247 bp long. This final contig was annotated using MITOS (Bernt et al., 2013). The locations of protein-coding genes were manually verified from the MITOS predictions by aligning to orthologous protein-coding gene sequences taken from the published mitochondrial genomes of lophotrochozoan taxa. Where possible, the locations of protein-coding genes were inferred to startb from the first in-frame start codon (ATN) and the C-terminal of the protein-coding genes inferred to be the first in-frame stop codon (TAA, TAG or TGA). Contigs from *Dicyema* sp. and *D. japonicum* identified as containing mitochondrial protein-coding genes were verified and annotated in the same way.

The secondary structure of tRNAs identified in the *I. linei* mitochondrial genome were inferred using the Mitfi program within MITOS. MITOS was also used to screen the *Dicyema* sp. and *D. japonicum* contigs which had been positively-identified as containing mitochondrial genes using blast. From MITOS, we could identify one reliable tRNA sequence for trnV on the same contig as *cox3* from *Dicyema* sp. Secondary structure for this tRNA was inferred using Mitfi in MITOS.

### Dicyema internal phylogeny

The *cox1* nucleotide sequences from *Dicyema* sp. and *D. japonicum* were aligned to publicly available *cox1* nucleotide sequences from other *Dicyema* species, and outgroup *cox1* sequences from diverse lophotrochozoan taxa (see Supplementary Table S2). Sequences were aligned using Muscle v.3.8.31 (Edgar, 2004) visualised in Mesquite v.3.31 (http://mesquiteproject.org) and trimmed to remove uninformative residues. Maximum Likelihood inference was carried out on the trimmed alignments using IQ-Tree (v.1.6.1) (Nguyen et al., 2015), letting the implemented model testing (ModelFinder (Kalyaanamoorthy et al., 2017)) pick the best-suited phylogenetic model (TVM+I+G4), and performing 1000 bootstrap replicates (UFBoot(Hoang et al., 2018)). The tree was visualised using Seaview v.4.4.2 (Gouy et al., 2010) and annotated with Inkscape.

### Data availability

All mitochondrial genome data presented in this study have been submitted to NCBI GenBank, under accession numbers as follows: *Intoshia linei* mitochondrial genome sequence, MG839537; *Dicyema* sp. genes, MG839520-MG839528; *Dicyema japonicum* genes, MG839529-MG839536.

## Results

### *Intoshia linei* mitochondrial genome composition

The final *I. linei* mitochondrial genome sequence we could reconstruct was 14,247 base pairs long, but we were unable to close the circular genome with paired end reads. This may be attributed to the missing sequence being an AT-rich repetitive region, making it difficult to resolve. The *Intoshia linei* mitochondrial genome is very AT rich (83.40% AT); overall nucleotide usage on the forward strand is A = 38.98%; T = 44.42%; G = 8.56% and C = 8.04. Absolute AT-skew is 0.065 and GC-skew is 0.031.

Using MITOS (Bernt et al., 2013) and manual verification, we were able to predict the full reading frames of 12 protein-coding genes, 20 tRNAs and the small subunit rRNA (*rrnS*). No sequences resembling *atp8, trnQ* or *trnR* could be found in the final sequence. It is possible that *rrnL* is found between *trnC* and *trnM* based on a weak prediction using MITOS, but this could not be confirmed by aligning to known *rrnL* sequences. Genes in the *I. linei* mitochondrial genome are found in two blocks of opposing transcriptional polarity: those in the first 3104 base pairs are found on the reverse strand (*trnK-trnV-trnT-trnY-nad4-trnS2-cox1-trnF*) and all other genes are found on the forward strand, where the forward strand is defined as that containing a greater portion of the protein-coding sequence (Fig. 1, Table 1). All protein-coding genes have standard initiation codons (ATA x 8; ATG x 1; and ATT x 3); ten of the protein-coding genes have a standard termination codon (TAA), with the exception of *nad5* and *nad6*, which appear to have a truncated stop codon (T— and TA-, respectively) (Table 1).

**Figure 1:**
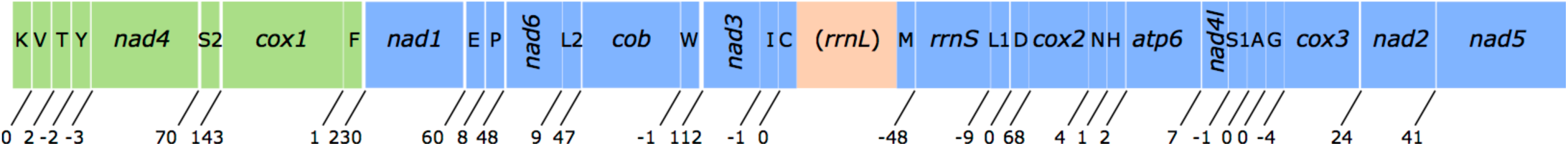
Overview of the mitochondrial sequence resolved for *Intoshia linei*. Genes not drawn to scale. Numbers beneath the sequences show intergenic spaces (positive values) or intergenic overlap (negative values). Protein-coding genes are denoted by three letter abbreviations; ribosomal genes by four letter abbreviations. tRNAs are shown by single uppercase letters (for recognised codons of L1, L2, S1 and S2 see Table 1). Genes found on the negative (reverse) strand are coloured green; genes found on the positive (forward) strand are coloured blue. The unreliable prediction for *rrnS* is shown in orange.

**Table 1:** Organisation of the *I. linei* mitochondrial genome. Uncertain position of *rrnL* shown in red.

| Feature | Strand | Start | Stop | Length<br>(nucleotides) | Length<br>(AA) | Start<br>Codon | Stop<br>Codon | Intergenic<br>region |
| --- | --- | --- | --- | --- | --- | --- | --- | --- |
| *trnK* | - | 99 | 157 | 59 |  |  |  | 0 |
| *trnV* | - | 158 | 209 | 52 |  |  |  | 2 |
| *trnT* | - | 212 | 271 | 60 |  |  |  | -2 |
| *trnY* | - | 270 | 330 | 61 |  |  |  | -3 |
| *nad4* | - | 328 | 1509 | 1182 | 394 | ATA | TAA | 70 |
| *trnS2* (TGA) | - | 1580 | 1637 | 58 |  |  |  | 143 |
| *cox1* | - | 1781 | 3175 | 1395 | 465 | ATA | TAA | 1 |
| *trnF* | - | 3177 | 3237 | 61 |  |  |  | 230 |
| *nad1* | + | 3468 | 4358 | 891 | 297 | ATG | TAA | 60 |
| *trnE* | + | 4419 | 4470 | 52 |  |  |  | 8 |
| *trnP* | + | 4479 | 4539 | 61 |  |  |  | 48 |
| *nad6* | + | 4588 | 5028 | 441 | 147 | ATT | TA- | 9 |
| *trnL2* (TAA) | + | 5038 | 5098 | 61 |  |  |  | 47 |
| *cob* | + | 5146 | 6168 | 1023 | 341 | ATA | TAA | -1 |
| *trnW* | + | 6168 | 6224 | 57 |  |  |  | 112 |
| *nad3* | + | 6337 | 6687 | 351 | 117 | ATA | TAA | -1 |
| *trnI* | + | 6687 | 6749 | 63 |  |  |  | 0 |
| *trnC* | + | 6750 | 6804 | 55 |  |  |  | -1 |
| (*rrnL*) | + | 6804 | 8031 | 1228 |  |  |  | 1 |
| *trnM* | + | 8033 | 8096 | 64 |  |  |  | -48 |
| *rrnS* | + | 8049 | 8790 | 742 |  |  |  | -9 |
| *trnL1* (TAG) | + | 8782 | 8838 | 57 |  |  |  | 0 |
| *trnD* | + | 8839 | 8894 | 56 |  |  |  | 68 |
| *cox2* | + | 8963 | 9580 | 618 | 206 | ATA | TAA | 4 |
| *trnN* | + | 9585 | 9649 | 65 |  |  |  | 1 |
| *trnH* | + | 9651 | 9705 | 55 |  |  |  | 2 |
| *atp6* | + | 9708 | 10394 | 687 | 229 | ATA | TAA | 7 |
| *nad4l* | + | 10402 | 10677 | 276 | 92 | ATT | TAA | -1 |
| *trnS1* (TCT) | + | 10677 | 10728 | 52 |  |  |  | 0 |
| *trnA* | + | 10729 | 10781 | 53 |  |  |  | 0 |
| *trnG* | + | 10782 | 10837 | 56 |  |  |  | -4 |
| *cox3* | + | 10834 | 11613 | 780 | 260 | ATA | TAA | 24 |
| *nad2* | + | 11638 | 12507 | 870 | 290 | ATT | TAA | 41 |
| *nad5* | + | 12549 | 14135 | 1587 | 529 | ATA | T-- |  |

In the final 14,247 base pair long sequence, protein-coding genes account for 70.90% of the sequence (allowing for overlap between genes); tRNAs 8.13%; rRNAs (including the uncertain prediction for rrnL) 13.83%; and non-coding DNA 6.85%. Four regions of non-coding sequence greater than 100 base pairs are found in the *I. linei* genome: a 143 bp-long region between *trnS2* and *cox1*; 230 bp between *trnF* and *nad1* (where the two genes are found on opposite strands); 112 bp between *trnW* and *nad3;* and 112 bp following the *nad5* at the end of the genomic sequence. There is very little overlap between coding sequences: *rrnS* and the best prediction for *trnM* location overlap by 48 nucleotides, and there are eight incidences of overlap of coding sequences of fewer than 10 nucleotides across the sequence.

20 out of 22 typical tRNAs were identified in the *I. linei* mitochondrial genome (Fig. 2). All predicted tRNAs have an amino-acyl acceptor stem composed of seven or eight base pairs and an anticodon stem composed of four or five base pairs, with the exception of *trnA* and *trnV*, which appear to have truncated acceptor stems. The structure of the DHU arm is, for the most part, consistent with standard tRNA secondary structure: 14 of the predicted tRNAs have a typical four or five base pair DHU stem; four tRNAs have a truncated or modified DHU arm (*trnI, trnL1, trnL2* and *trnP*); and two tRNAs appear to have lost the DHU arm entirely (*trnS1* and *trnS2*). More unusually, the TC arm in almost all of the predicted tRNAs is either truncated or replaced by a TV loop. The only tRNA found to have a ‘typical’ TC arm structure is *trnS2*.

**Figure 2:**
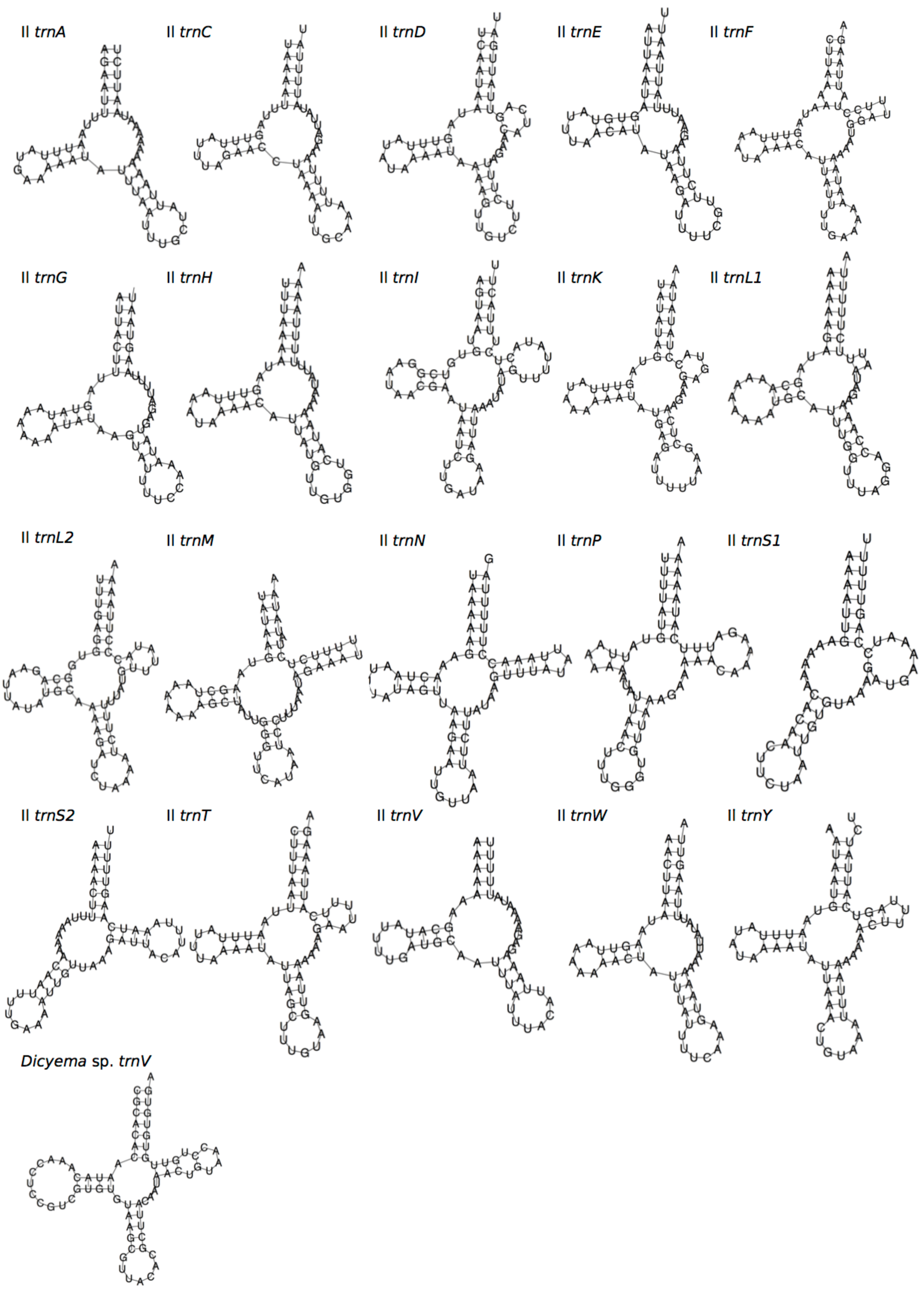
Predicted secondary structures of tRNAs from the mitochondrial genome sequence of *Intoshia linei* (annotated as *Il*), and of *trnV* from a putative mitochondrial contig of *Dicyema* sp. All secondary structures predicted by MiTFi in Mitos^28^.

### Gene order

Using CREx (Bernt et al., 2007), the protein-coding and rRNA gene order of *I. linei* was compared to a number of other metazoan taxa, including representatives from across the Lophotrochozoa, Ecdysozoa and Deuterostomia (see Supplementary Table S1). CREx analysis aims to identify common gene intervals between different mitochondrial genomes, and infer the reversals, transpositions, and reverse transpositions required to obtain a given gene order from the mitochondrial gene orders of other species. Compared with the other taxa included for analysis, *I. linei* has a highly divergent mitochondrial gene order, with the closest similarity found to the mitochondrial genome of the carmine spider mite *Tetranychus cinnabarinus* (Arthropoda; Chelicerata) (Fig.3). However, similarity in gene order between these genomes is still low, with the two sharing just four short conserved gene ‘blocks’, and with variation in gene order even within these common intervals. Of the lophotrochozoan species included for analysis, the species with the highest similarity to *I. linei* was found to be the nemertean *Paranemertes peregrina* (Fig. 3). Both species share the common arrangement of *nad1-nad6-cob*, and the adjacency of *rrnL-rrnS-cox2-atp6* and *nad4-nad5*, but with variation in the order of these two blocks. As with the similarity to *T. cinnabarinus*, conservation of gene order of *I. linei* compared to *P. peregrina* is very low given the number of possible shared gene boundaries. It is therefore evident that the gene order of *I. linei* is novel and very divergent amongst published metazoan mitochondrial genomes.

**Figure 3:**
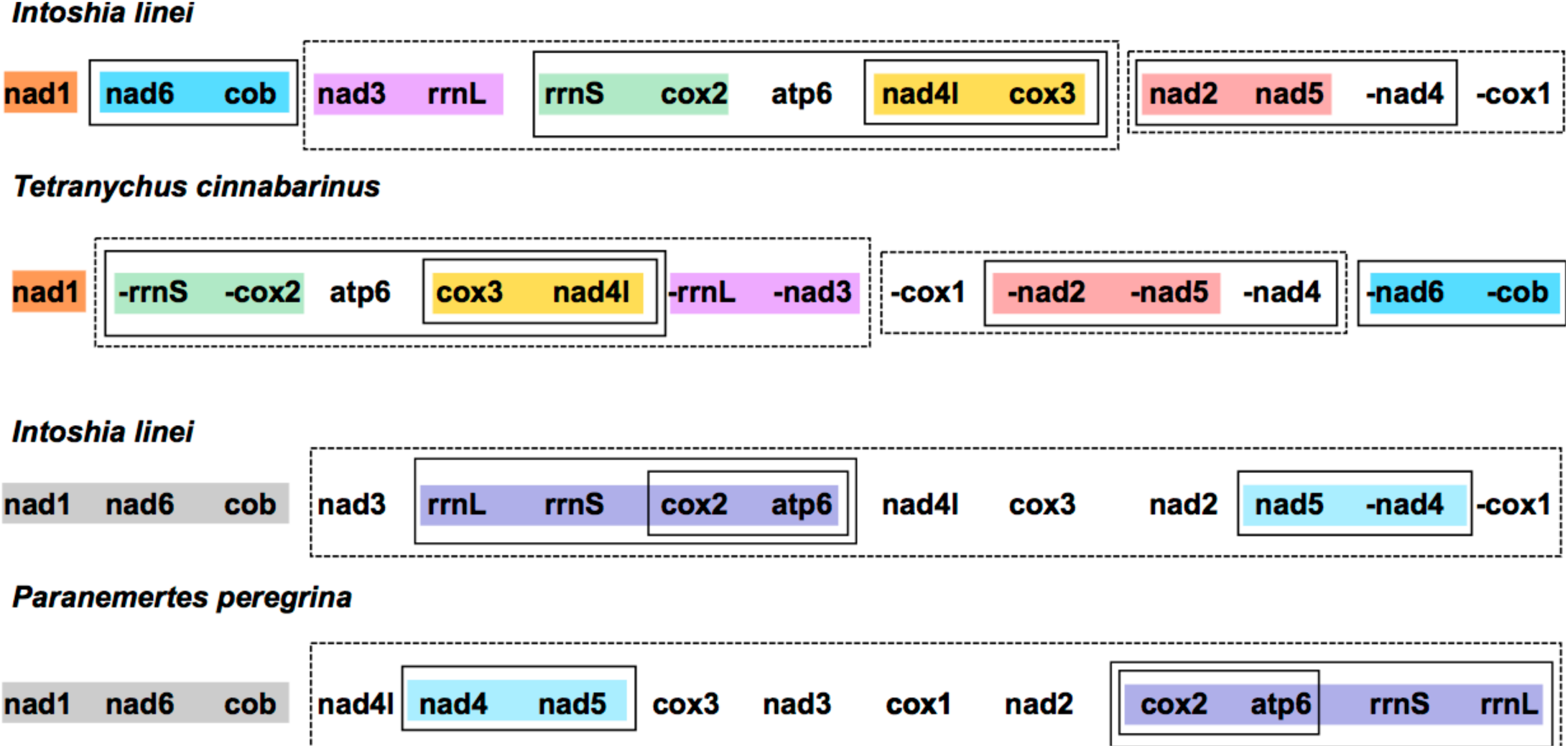
Gene order comparison between the mitochondrial genomes of *Intoshia linei*, the chelicerate *Teteranychus cinnabarinus* and the nemertean *Paranemertes peregrina*, as analysed using CREx. Genes on the minus strand are denoted by a minus sign (-X). Conserved blocks of genes between *I. linei* and *T. cinnabarinus* are shown in orange (*nad1*), blue (*nad6, cob*), pink (*nad3, rrnL*), green (*rrnS, cox2*), yellow (*nad4l, cox3*) and red (*nad2, nad5*). Variation in gene order and translocation between strands is found within these blocks between the two species. Conserved blocks of genes between *I. linei* and *P. peregrina* are shown in grey (*nad1, nad6, cob*), purple (*rrnL, rrnS, cox2, apt6*) and light blue (*nad4, nad5*). As before, variation in gene order and translocation between strands is evident between these common intervals. For both comparisons, solid or dashed-line boxes show larger regions of the genomes that encompass the same genes, but with no conserved gene order.

### *Dicyema sp.* and *Dicyema japonicum* mitochondrial genes

Using blast and manual sequence verification of contigs assembled using Trinity (Haas et al., 2013) and the CLC assembly cell, we were able to identify nine reconstructed mitochondrial transcripts containing protein-coding genes from *Dicyema* sp. (*cox1, 2, 3*; *cob*; and *nad1, 2, 3, 4* and *5*). In *D. japonicum* we found contigs for eight mitochondrial transcripts of protein-coding genes (*cox1, 2, 3*; *nad1, 3, 4* and *5*; and *cob*). The identification of *cob* and *nad3, 4* and *5* are novel for this taxon. All of the complete *Dicyema* protein-coding genes identified have full initiation and termination codons, with the exception of *D. japonicum nad1*, which has a TA-truncated stop codon. For each instance of a successfully identified gene-bearing contig from *Dicyema* sp., we also found additional contigs that had strongly matching blast hits to *Octopus* or other cephalopods, and thus are most likely contamination from the host species.

All mitochondrial protein-coding genes found for *Dicyema* were located on individual contigs. In no instance were two or more protein-coding genes found on the same reconstructed transcript. However, each reconstructed mitochondrial transcript did contain non-coding sequence in addition to protein-coding gene sequence (Fig. 4). The length and location (5’ and/or 3’) of non-coding sequence in the reconstructed transcripts was variable between species and between genes (Fig. 4). For each protein-coding gene, we compared contigs assembled using both the CLC-assembler and Trinity in order to identify reconstructed mitochondrial transcripts with the longest stretch of protein-coding sequence. Of the 17 dicyemid genes we report, seven were derived from CLC-assembled contigs (*cox1, 2, 3, nad3, 5* from *D. japonicum*, and *nad2, nad5* from *Dicyema* sp.), and ten from Trinity-assembled contigs (*cob, nad1, 4* from *D. japonicum* and *cox1, 2, 3, cob, nad1, 3, 4* from *Dicyema* sp.). In addition, the best reconstructed mitochondrial contig we identified for *nad1* (*Dicyema* sp.) was found to contain duplicated stretches of identical protein-coding gene sequence. We attribute this to an assembly artefact rather than speculating about potential evidence for mini-circles. We also identified a frameshift in the coding sequence for *nad2* from *Dicyema* sp. in the longest reconstructed mitochondrial contig from this species (Fig. 4).

**Figure 4:**
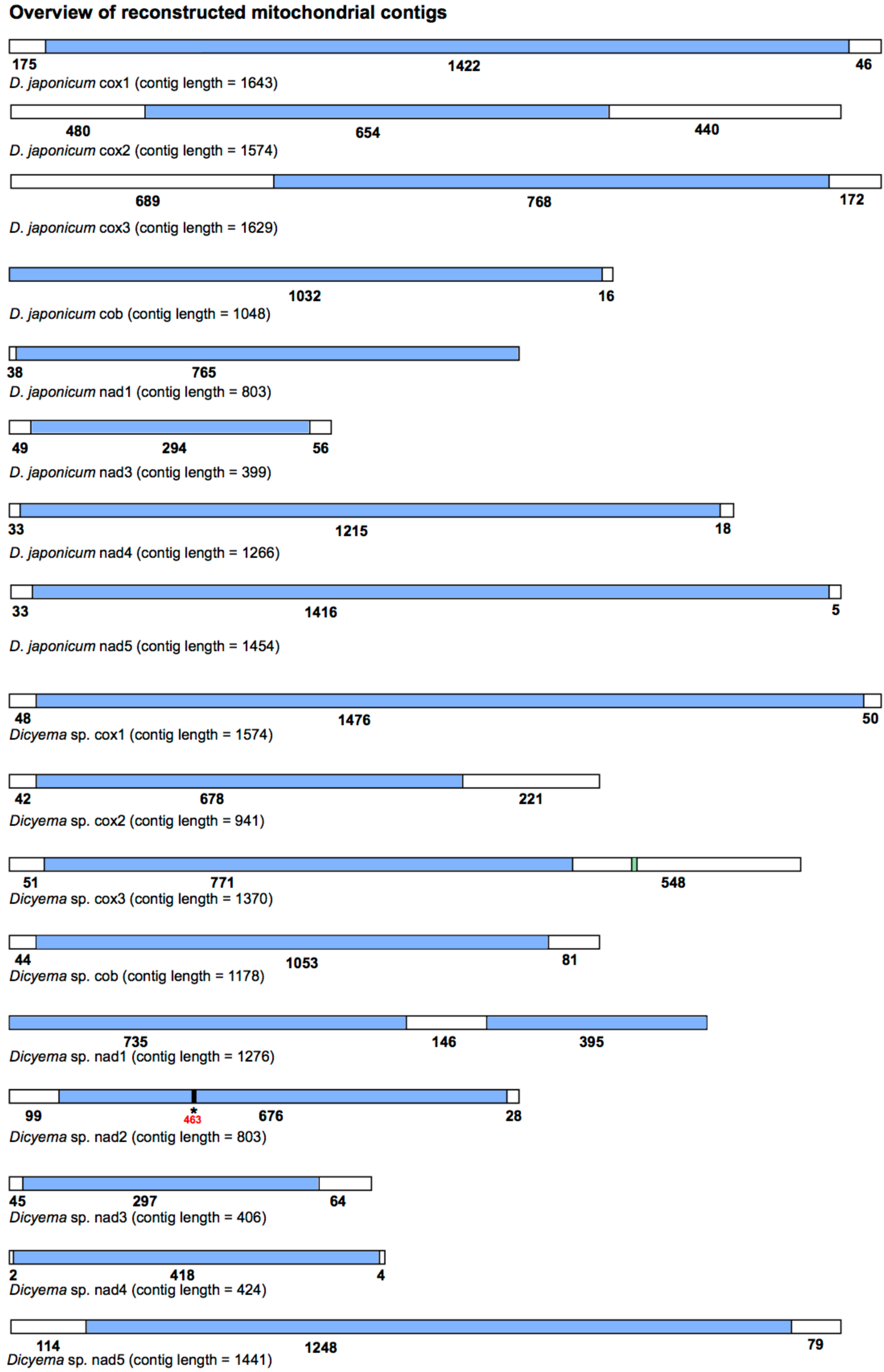
Overview of the reconstructed mitochondrial transcripts of protein-coding genes found in *Dicyema japonicum* and *Dicyema* sp. Transcripts are not to scale. Regions coloured blue indicate protein-coding sequence; regions coloured white indicate non-coding sequence. Numbers under each transcript correspond to the length of each respective coding or non-coding region, in base pairs. For *cox3* in *Dicyema* sp. the location of the putative *trnV* sequence is shown in green (nucleotides 1130-1200). For *nad2* in *Dicyema* sp., the location of a frameshift at position 463 in the longest coding sequence assembly is denoted by an asterisk (^*^)

Screening the *Dicyema* sp. gene-containing contigs using MITOS standalone software, we were also able to predict one reliable tRNA sequence for *trnV* adjacent to *cox3* (Fig. 4). *Dicyema sp. trnV* has an eight bp acceptor stem; a five bp anti-codon stem; and a four base pairs DHU stem (Fig. 2). As was found in *I. linei*, the TC arm appears to be modified from a standard ‘cloverleaf’ structure. In no other cases did we find any sequence from more than one gene on a single contig.

### Dicyema internal phylogeny using *cox1*

*Cox1* is commonly used as a species ‘barcoding’ gene, and can be used in phylogenetic inference to discriminate between closely related species (Hebert et al., 2003). Given that a number of *cox1* genes have been sequenced from different *Dicyema* species, we aimed to use publicly available *Dicyema cox1* sequences along with the two new *cox1* sequences found in this study for phylogenetic inference, and to determine the relationship of *Dicyema* sp. And *D. japonicum* to other dicyemids. In aligning the *cox1* nucleotide sequences from *Dicyema* members to other metazoans, we found that all *Dicyema cox1* sequences had several conserved deletions, not present in other metazoan *cox1* sequences included in alignment (Fig. 5). These comprise in-frame deletions of two, five, four and two amino acids moving from the N-terminus to the C-terminus of the protein. These deletions appear to be unique to members of the *Dicyema*, and were present in all *Dicyema* species included in alignment.

**Figure 5:**
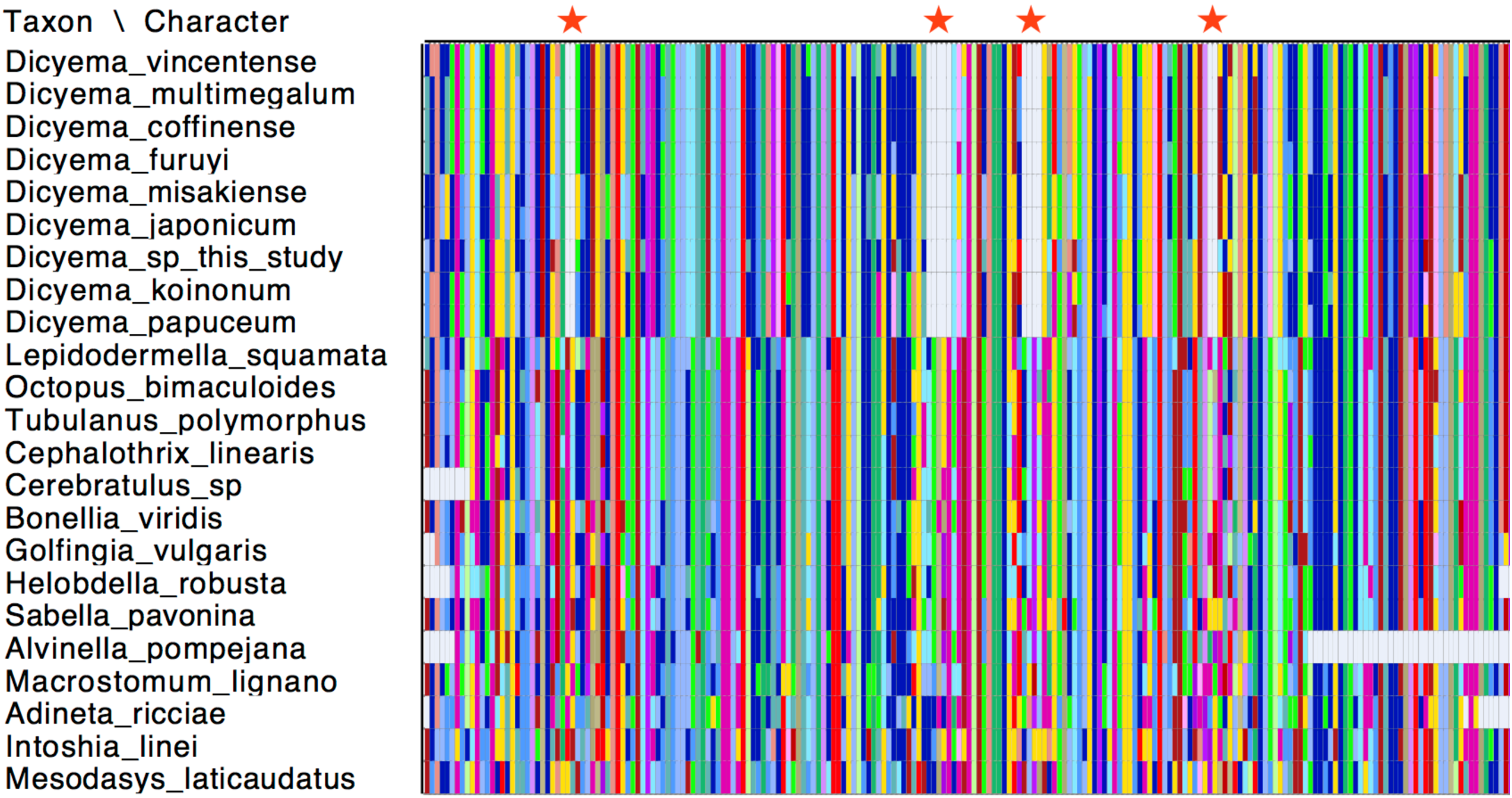
Conserved deletions in the amino acid sequence of *cox1* taken from publicly available dicyemid sequences (*D. vincentense, D. multimegalum, D. coffinense, D. furuyi, D. misakiense, D. koinonum, D. papuceum*) and the *Dicyema* species presented in this analysis (*D. japonicum* and *Dicyema* sp.), in alignment with other lophotrochozoan *cox1* sequences. The location of the deletions (two amino acids; five amino acids; four amino acids and two amino acids) moving from the N-terminus to the C-terminus of the protein, are shown by red stars. Colours in the alignment correspond to amino acid colours as used by Mesquite v.3.31.

Maximum likelihood phylogenetic analysis was carried out using *cox1* sequences from publicly available *Dicyema* species; the *D. japonicum* and *Dicyema* sp. *cox1* sequences assembled in this study; and *cox1* sequences from a diverse representation of lophotrochozoans as outgroup taxa (Supplementary Table S2). As anticipated, the dicyemids included in analysis form their own branch on the tree. This analysis found a close affinity for *D. japonicum* with *D. misakiensi*, with 97% bootstrap support at this node. At the nucleotide level, sequences from *D. japonicum* and *D. misakiensi* are ∼98% identical. The remaining *Dicyema* species branch separately from *D. japonicum* and *D. misakiense*, with the *Dicyema* sp. sequence from our analysis branching separately from the other six representatives of the group. As found for *D. japonicum* and *D. misakiense*, the *cox1* sequences from *D. multimegalum* and *D. coffinense* are ∼98% identical at the nucleotide level. (Fig. 6)

**Figure 6:**
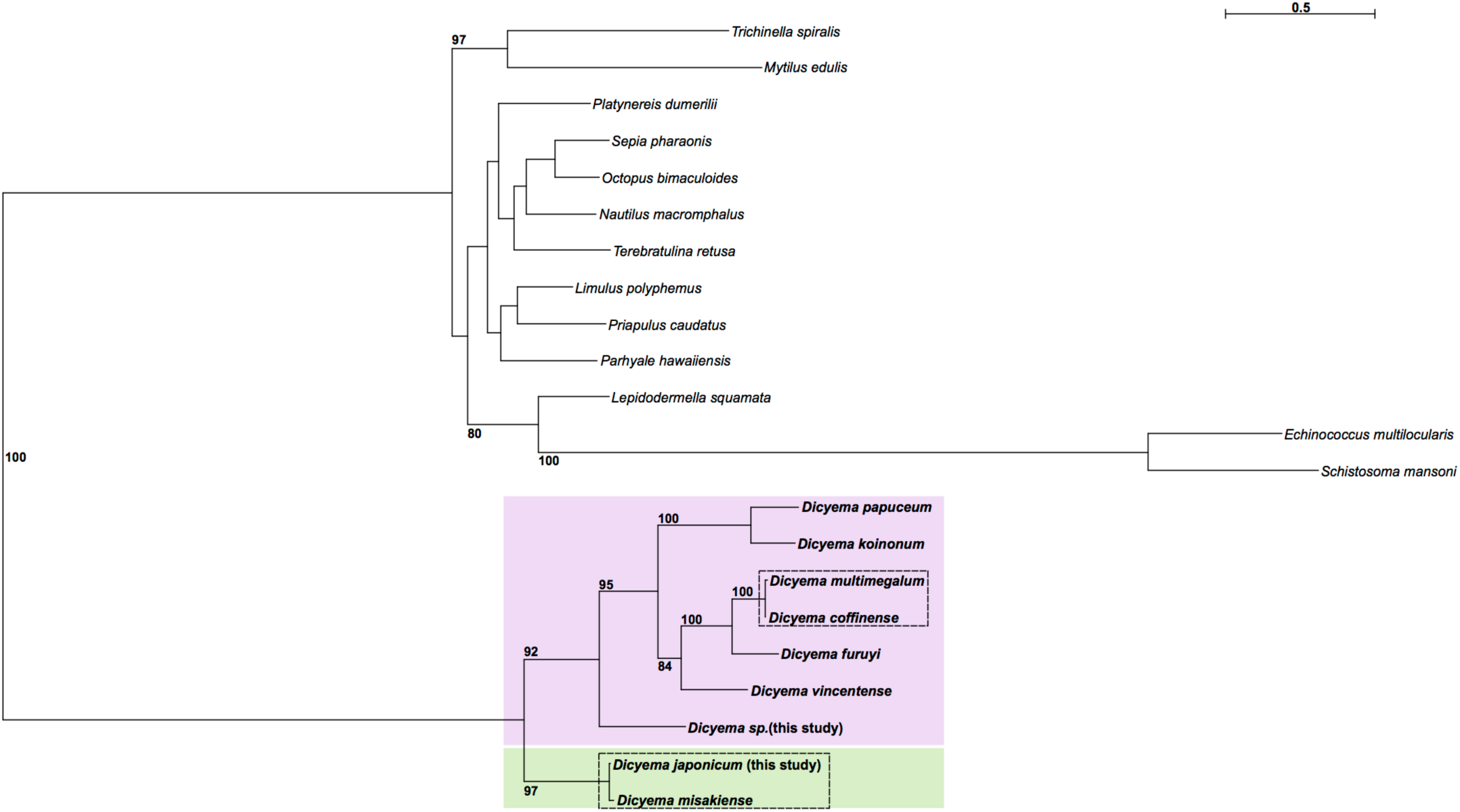
Phylogenetic analysis of *cox1* genes from dicyemids using Maximum Likelihood (RAxML^30^), including the two *Dicyema* species presented in this analysis (*Dicyema* sp. and *D. japonicum*). *Cox1* sequences from diverse lophotrochozoan taxa included as outgroups. Bootstrap support is shown at relevant nodes. Phylogenetic inference shows a split of dicyemid species into two groups, one containing *D. japonicum* and *D. misakiense* (green box), and the other containing all other dicyemid species included in analysis (pink box). Within the second group, *Dicyema* sp. is found on a separate branch from the rest of the included dicyemids. Very short branch lengths between both *D. coffinense* and *D. multimegalum*, and *D. japonicum* and *D. misakiense*, indicate that they are very closely related or could be identical species (dashed-line boxes).

## Discussion

Although the mitochondrial genome we found from *I. linei* is not demonstrated to be a closed circular molecule, this analysis presents the first mitochondrial genome from an orthonectid species, and includes 12 protein-coding genes, 20 tRNAs and both ribosomal RNAs. *Atp8* was not found, but this gene has been lost from the mitochondrial genomes of taxa in many different metazoan lineages (Boore, 1999) and so its absence from our assembly may be real rather than representing missing data. Compared to the drastically reduced *I. linei* nuclear genome, its mitochondrial genome has a gene complement that is fairly standard across the Metazoa(Mikhailov et al., 2016). Genes in the *I. linei* mitochondrial genome are clustered into two different blocks of opposite transcriptional polarity. The block comprising *trnK-trnV-trnT-trnY-nad4-trnS2-cox1-trnF* at the ‘start’ of the sequence are found on the negative strand, whilst all other genes are transcribed from the positive strand, suggesting an inversion event (Fig. 1).

The ∼84% A+T content found in the *I. linei* mitochondrial genome is high compared to other invertebrate mitochondrial genomes – for example, the chelicerate *Limulus polyphemus* (A+T content = 67.6%) (Lavrov et al., 2000) and the annelid *Lumbricus terrestris* (A+T content = 61.6%) (Boore & Brown, 1995). The very high A+T content of the mitochondrial genome is even higher than the high nuclear genome A+T content of *I. linei* (73%), and coincides with a very fast rate of mitochondrial evolution in this species.

The small proportion of non-coding mtDNA we found for *I. linei* is typically true of mitochondrial genomes. Although there is very little gene overlap in the sequence found for *I. linei*, the sequence for *trnM* is predicted with significant overlap with *rrnS*. Whilst it is possible that this is a computational mis-prediction, large gene overlaps have been reported in other mitochondrial genomes, and this overlap could be an approach to reduce mitochondrial genome size (Robertson et al., 2017).

Gene order in the *I. linei* mitochondrial genome is highly divergent from the gene order that is commonly found in other Metazoa. Gene order in the mitochondrial genomes of different lineages are largely stable, with the rearrangement of protein-coding genes occurring relatively infrequently. Where rearrangement events do occur, they are thought to be a result of tandem duplications and multiple random deletions. In this model, a portion of the mitochondrial genome is erroneously duplicated, and the subsequent random loss of one copy of a gene (by deletion or the accumulation of mutations) results in a novel gene order (Boore, 2000).

Our analysis demonstrated that *I. linei* has a highly divergent mitochondrial gene order in comparison to other (published) metazoan mitochondrial genomes. Of the species included – chosen as a broad representation of different metazoan lineages (see Supplementary Table S1) – the closest similarity was found to be to the chelicerate *T. cinnabarinus* (Fig. 3). However, of the highest possible conserved gene order score of 204 (that is, when two mitochondrial genomes have identical gene orders for the 12 protein-coding genes (*atp8* is not present in *I. linei*) and two rRNAs included in the CREx matrix calculation), similarity of gene order between *I. linei* and *T. cinnabarinus* was still comparatively poor, at just 32. In accordance with the proposed affinity of *I. linei* to the Lophotrochozoa, the highest-scoring similarity to the lophotrochozoan species included in analysis was with the nemertean *P. peregrina*. Again, the two species share comparatively little gene order conservation: just three common gene blocks were identified, one of which (*rrnL-rrnS-cox2-atp6*) has further rearrangement therein (Fig. 3). Of the common intervals identified, the block of *nad1-nad6-cob* is conserved between *I. linei* and *P. peregrina*, and it is possible that this arrangement is plesiomorphic within the Lophotrochozoa.

Interestingly, analysis of five mitochondrial genomes from early branching annelids found that gene order varied greatly between these species(Weigert et al., 2016), and other studies of various lophotrochozoan mitochondrial genomes – including Brachiopoda (*Lingula anatina*) (Luo et al., 2015); various *Schistosoma* species (Webster & Littlewood, 2012); and nemerteans (Podsiadlowski et al., 2009) – indicate that extensive gene order rearrangements have occurred in different lophotrochozoan lineages. It is likely that the divergent gene order in *Intoshia linei* can be associated with the parasitic lifestyle and rapid rate of evolution seen for this species: studies in a number of other parasitic taxa indicate that an accelerated rate of gene rearrangement in mitochondrial genomes could be associated with this lifestyle. For example, various *Schistosoma* species have mitochondrial genomes with a unique gene order (Le et al., 2000), and this has also been observed in the ectoparasitic louse *Heterodoxus macropus* (Shao et al., 2001), various species of mosquito (Beard et al., 1993), and parasitic hymenopterans (Dowton & Austin, 1999), amongst others.

All of the tRNAs predicted from *I. linei* and the one tRNA found in *Dicyema sp.* have deviations from the ‘standard’ secondary structure of the TC arm (Fig. 2). Although it is unusual amongst typical metazoan mitochondrial genomes to have such consistent modifications to one element of the tRNA cloverleaf structure, a great deal of variation can be found in tRNA structures across the Metazoa. Mitochondrial genomes with almost all tRNAs lacking either the TC arm or DHU arm – termed ‘minimal functional tRNAs’ – have been reported in a number of different lineages. In nematodes, analysis of tRNAs with TV loops in place of a TC arm suggests that an ‘L-shaped’ tRNA, analogous to a typical cloverleaf-structure tRNA – can maintain normal tertiary interactions and remain functional. Furthermore, it is likely that the tRNAs reported in this analysis are functional, as the acceptor stems and anti-codon stems are, for the most part, complete, and it is highly likely that they would have accumulated mutations should they have lost functionality. Instead, the reduction we observe in tRNA secondary structure could be a result of selective pressure to reduce the TC arm, and provide another example of minimally functional tRNAs, in addition to those already found across the Metazoa.

Previous analyses had suggested that mitochondrial genes (*cox1, cox2* and *cox3*) in dicyemids are found on individual mini-circles in somatic cells, as opposed to being located on a larger circular mitochondrial genome (Catalano et al., 2015; Watanabe et al., 1999).

Mitochondrial mini-circles – although rare across the Metazoa - do appear to be most prevalent in parasitic species. A number of studies have reported the presence of mini-circle mitochondrial genomes, fragmented to various degrees, in a number of lice and nematode species (Cameron et al., 2011; Dong et al., 2014; Hunt et al., 2016; Phillips et al., 2016; Shao et al., 2009).

We identified a mitochondrial contig for *nad1* in *Dicyema* sp. that contained repeated *nad1* protein-coding sequence. This could be an assembly artefact induced by a circular *nad1* molecule, but the question of the presence of minicircles remains unresolved in our analysis. Further investigating the validity of mitochondrial mini-circles was outside of the scope of the present study, but future approaches involving long-range PCR or long-read sequencing should be conducted to resolve this question.

All dicyemid *cox1* sequences were found to have a series of four in-frame deletions within a region of the gene that was well-aligned with *cox1* sequences taken from other invertebrate species (Fig. 5). Insertions and deletions (indels) in genes are rare genomic changes that can be used to infer common evolutionary history(Belinky et al., 2010), but this set of conserved deletions are so far known only from the *Dicyema* genus. No such deletions are found in the *I. linei cox1* gene, or in the same protein-coding sequence taken from across the Lophotrochozoa.

The internal phylogeny of dicyemids was inferred using Maximum Likelihood based on *cox1* gene sequences. Our analysis found a split of the dicyemids into two groups: one comprising *D. japonicum* and *D. misakiense*, and the other comprising the seven other Dicyema species (Fig. 6). *D. coffinense* and *D. multimegalum* are likely to be very closely related or identical species, with both dicyemids isolated from Australian *Sepia* species (Catalano, 2013), and with a greater than 98% sequence similarity at the nucleotide sequence level. Our analysis also found a very close affinity for *D. japonicum* with *D. misakiensi*, with 97% bootstrap support at this node. As found for *D. coffinense* and *D*. *multimegalum* these *cox1* sequences are >98% identical at the nucleotide level, and it is thus likely that they are very closely related or the same species, both found in octopus living in the West North Pacific off the coast of Honshu Island, Japan. The *Dicyema* sp. *cox 1* assembled in this analysis also groups with these *Dicyema* members, despite being isolated from *Sepia* living off the coast of Florida, potentially hinting at long-range dispersal through the host species. Further investigation into dicyemid members isolated from hosts in other geographical locations could help to inform how phylogenetic structure of the parasitic *Dicyema* is reflective of dispersal of the host octopus (Tobias et al., 2017).

## Supplementary Information: The mitochondrial genomes of the mesozoans *Intoshia linei, Dicyema* sp., and *Dicyema japonicum*

Helen. E. Robertson, Philipp. H. Schiffer and Maximilian. J. Telford

**Supplementary Table S1:**
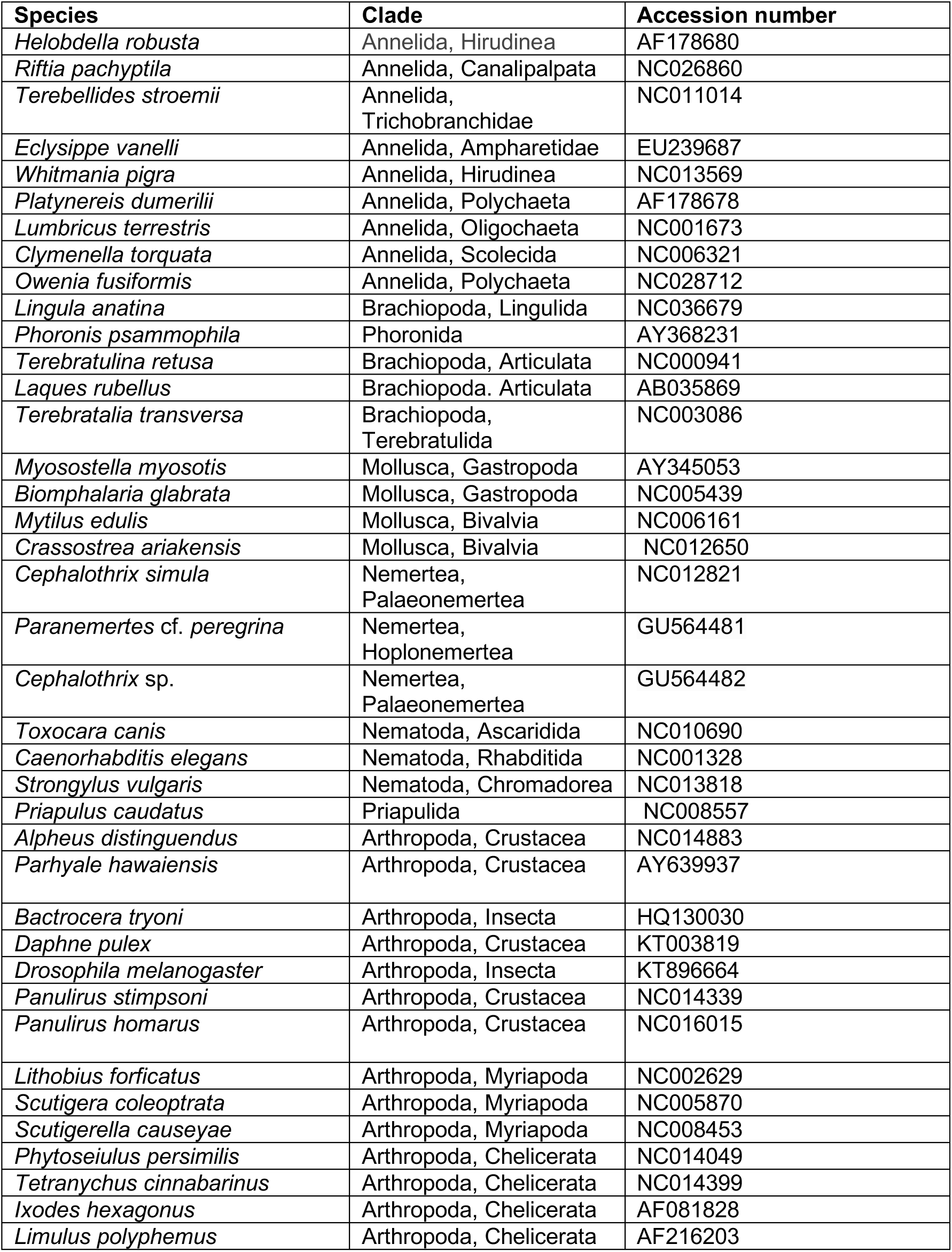
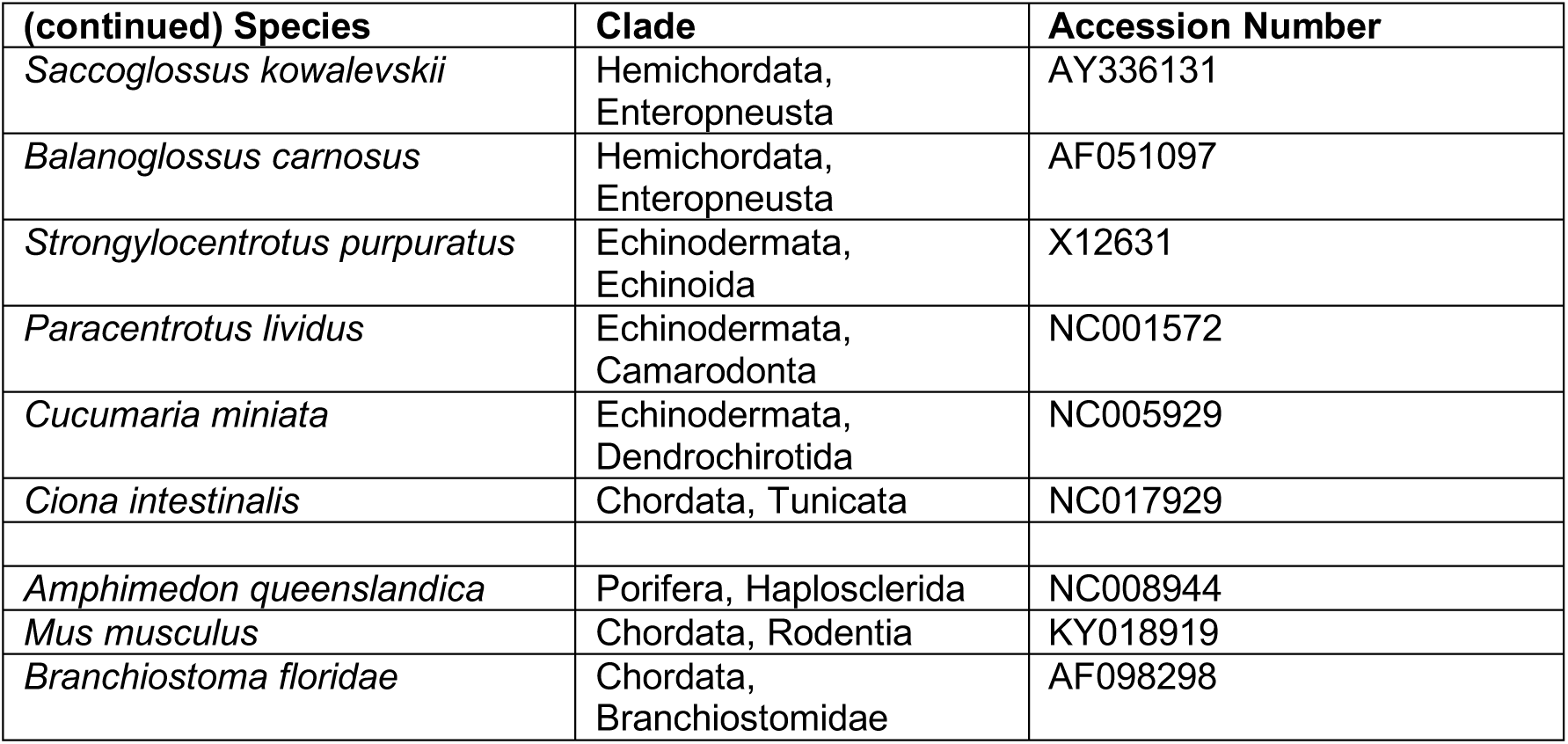
Names and Accession numbers (NCBI) of taxa used in CREx mitochondrial gene order comparison

**Supplementary Table S2:** Names and Accession numbers (NCBI) of taxa used in *cox1* phylogenetic analysis

| **Species** | **Clade** | **Accession** |
| --- | --- | --- |
| *Aplysia californica* | Mollusca, Gastropoda | YP003687 |
| *Lepidodermella squamata* | Platyzoa, Gastrotricha | NC026985 |
| *Echinococcus multilocularis* | Platyhelminthes,<br>Cyclophyllidea | AB385610 |
| *Schistosoma mansoni* | Platyhelminthes,<br>Diplostomida | AF216698 |
| *Sepia pharaonis* | Mollusca, Cephalopoda | KC632521 |
| *Octopus bimaculoides* | Mollusca, Cephalopoda | NC029723 |
| *Nautilus macromphalus* | Mollusca, Cephalopoda | DQ472026 |
| *Terebratulina retusa* | Brachiopoda, Terebratulida | NC000941 |
| *Platynereis dumerilii* | Annelida, Polychaeta | AF178678 |
| *Limulus polyphemus* | Arthropoda, Chelicerata | AF216203 |
| *Priapulus caudatus* | Priapulida | NC008557 |
| *Parhyale hawaiiensis* | Arthropoda, Crustacea | AY639937 |
| *Trichinella spiralis* | Nematoda, Trichocephalida | KM357422 |
| *Mytilus edulis* | Mollusca, Bivalvia | NC006161 |
| *Dicyema papuceum* | Dicyemida | KF208352 |
| *Dicyema koinonum* | Dicyemida | KF208346 |
| *Dicyema multimegalum* | Dicyemida | KF208351 |
| *Dicyema coffinense* | Dicyemida | KF208317 |
| *Dicyema furuyi* | Dicyemida | KF208321 |
| *Dicyema vincentense* | Dicyemida | KF208356 |
| *Dicyema misakiense* | Dicyemida | AB011832 |

## References

Awata, H., Noto, T. and Endoh, H. (2005). Differentiation of somatic mitochondria and the structural changes in mtDNA during development of the dicyemid *Dicyema japonicum* (Mesozoa). Molecular Genetics and Genomics, 273, 441–449. doi:10.1007/s00438-005-1157-2.

Beard, C. B., Hamm, D. M. and Collins, F. H. (1993). The mitochondrial genome of the mosquito Anopheles gambiae: DNA sequence, genome organization, and comparisons with mitochondrial sequences of other insects. Insect Molecular Biology, 2, 103–124. doi:10.1111/j.1365-2583.1993.tb00131.x.

Belinky, F., Cohen, O. and Huchon, D. (2010). Large-Scale Parsimony Analysis of Metazoan Indels in Protein-Coding Genes. Molecular Biology and Evolution, 27, 441–451. doi:10.1093/molbev/msp263.

Bernt, M., Donath, A., Jühling, F., Externbrink, F., Florentz, C., Fritzsch, G., Pütz, J., Middendorf, M. and Stadler, P. F. (2013). MITOS: improved de novo metazoan mitochondrial genome annotation. Molecular Phylogenetics and Evolution, 69, 313–319. doi:10.1016/j.ympev.2012.08.023.

Bernt, M., Merkle, D., Ramsch, K., Fritzsch, G., Perseke, M., Bernhard, D., Schlegel, M., Stadler, P. F. and Middendorf, M. (2007). CREx: inferring genomic rearrangements based on common intervals. Bioinformatics, 23, 2957–2958. doi:10.1093/bioinformatics/btm468.

Bolger, A. M., Lohse, M. and Usadel, B. (2014). Trimmomatic: a flexible trimmer for Illumina sequence data. Bioinformatics, 30. doi:10.1093/bioinformatics/btu170.

Boore, J. L. (1999). Animal mitochondrial genomes. Nucleic Acids Res, 27, 1767–1780.

Boore, J. L. (2000). The Duplication/Random Loss Model for Gene Rearrangement Exemplified by Mitochondrial Genomes of Deuterostome Animals. In Comparative Genomics: Empirical and Analytical Approaches to Gene Order Dynamics, Map Alignment and the Evolution of Gene Families (eds. Sankoff, D., and Nadeau, J. H.), pp. 133–147. Springer Netherlands, Dordrecht.

Boore, J. L. and Brown, W. M. (1995). Complete sequence of the mitochondrial DNA of the annelid worm *Lumbricus terrestris*. Genetics, 141.

Boore, J. L., Lavrov, D. V. and Brown, W. M. (1998). Gene translocation links insects and crustaceans. Nature, 392, 667–668. doi:10.1038/33577.

Cameron, S. L., Yoshizawa, K., Mizukoshi, A., Whiting, M. F. and Johnson, K. P. (2011). Mitochondrial genome deletions and minicircles are common in lice (Insecta: Phthiraptera). BMC Genomics, 12, 394–394. doi:10.1186/1471-2164-12-394.

Catalano, S. R. (2013). Five new species of dicyemid mesozoans (Dicyemida: Dicyemidae) from two Australian cuttlefish species, with comments on dicyemid fauna composition. Systematic Parasitology, 86, 125–151. doi:10.1007/s11230-013-9443-6.

Catalano, S. R., Whittington, I. D., Donnellan, S. C., Bertozzi, T. and Gillanders, B. M. (2015). First comparative insight into the architecture of COI mitochondrial minicircle molecules of dicyemids reveals marked inter-species variation. Parasitology, 142, 1066–1079. doi:10.1017/S0031182015000384.

Dodson, E. O. (1956). A Note on the Systematic Position of the Mesozoa. Systematic Zoology, 5, 37–40. doi:10.2307/2411651.

Dong, W.-G., Song, S., Guo, X.-G., Jin, D.-C., Yang, Q., Barker, S. C. and Shao, R. (2014). Fragmented mitochondrial genomes are present in both major clades of the blood-sucking lice (suborder Anoplura): evidence from two Hoplopleura rodent lice (family Hoplopleuridae). BMC Genomics, 15, 751. doi:10.1186/1471-2164-15-751.

Dowton, M. and Austin, A. D. (1999). Evolutionary dynamics of a mitochondrial rearrangement ‘hotspot’ in the Hymenoptera.

Edgar, R. C. (2004). MUSCLE: multiple sequence alignment with high accuracy and high throughput. Nucleic Acids Research, 32. doi:10.1093/nar/gkh340.

Furuya, H. and Tsuneki, K. (2003). Biology of dicyemid mesozoans. Zool Sci, 20. doi:10.2108/zsj.20.519.

Gouy, M., Guindon, S. and Gascuel, O. (2010). SeaView Version 4: A Multiplatform Graphical User Interface for Sequence Alignment and Phylogenetic Tree Building. Molecular Biology and Evolution, 27, 221–224. doi:10.1093/molbev/msp259.

Haas, B. J., Papanicolaou, A., Yassour, M., Grabherr, M., Blood, P. D. and Bowden, J. (2013). De novo transcript sequence reconstruction from RNA-seq using the Trinity platform for reference generation and analysis. Nat Protoc, 8. doi:10.1038/nprot.2013.084.

Hebert, P. D. N., Ratnasingham, S. and deWaard, J. R. (2003). Barcoding animal life: cytochrome c oxidase subunit 1 divergences among closely related species. Proceedings of the Royal Society B: Biological Sciences, 270, S96–S99. doi:10.1098/rsbl.2003.0025.

Hoang, D. T., Chernomor, O., von Haeseler, A., Minh, B. Q. and Vinh, L. S. (2018). UFBoot2: Improving the Ultrafast Bootstrap Approximation. Molecular Biology and Evolution, 35, 518–522. doi:10.1093/molbev/msx281.

Hunt, V. L., Tsai, I. J., Coghlan, A., Reid, A. J., Holroyd, N., Foth, B. J., Tracey, A., Cotton, J. A., Stanley, E. J., Beasley, H., Bennett, H. M., Brooks, K., Harsha, B., Kajitani, R., Kulkarni, A., Harbecke, D., Nagayasu, E., Nichol, S., Ogura, Y., Quail, M. A., Randle, N., Xia, D., Brattig, N. W., Soblik, H., Ribeiro, D. M., Sanchez-Flores, A., Hayashi, T., Itoh, T., Denver, D. R., Grant, W., Stoltzfus, J. D., Lok, J. B., Murayama, H., Wastling, J., Streit, A., Kikuchi, T., Viney, M. and Berriman, M. (2016). The genomic basis of parasitism in the Strongyloides clade of nematodes. Nat Genet, 48, 299–307. doi:10.1038/ng.3495

Kalyaanamoorthy, S., Minh, B. Q., Wong, T. K. F., von Haeseler, A. and Jermiin, L. S. (2017). ModelFinder: fast model selection for accurate phylogenetic estimates. Nature Methods, 14, 587. doi:10.1038/nmeth.4285

Lavrov, D. V., Boore, J. L. and Brown, W. M. (2000). The Complete Mitochondrial DNA Sequence of the Horseshoe Crab Limulus polyphemus. Molecular Biology and Evolution, 17, 813–824. doi:10.1093/oxfordjournals.molbev.a026360.

Le, T. H., Blair, D., Agatsuma, T., Humair, P.-F., Campbell, N. J. H., Iwagami, M., Littlewood, D. T. J., Peacock, B., Johnston, D. A., Bartley, J., Rollinson, D., Herniou, E. A., Zarlenga, D. S. and McManus, D. P. (2000). Phylogenies Inferred from Mitochondrial Gene Orders—A Cautionary Tale from the Parasitic Flatworms. Molecular Biology and Evolution, 17, 1123–1125. doi:10.1093/oxfordjournals.molbev.a026393.

Lu, T.-M., Kanda, M., Satoh, N. and Furuya, H. (2017). The phylogenetic position of dicyemid mesozoans offers insights into spiralian evolution. Zoological Letters, 3, 6. doi:10.1186/s40851-017-0068-5.

Luo, Y.-J., Satoh, N. and Endo, K. (2015). Mitochondrial gene order variation in the brachiopod *Lingula anatina* and its implications for mitochondrial evolution in lophotrochozoans. Marine Genomics, 24, Part 1, 31–40. doi:https://doi.org/10.1016/j.margen.2015.08.005.

Margulis, L. and Chapman, M. J. (2009). Chapter Three - ANIMALIA. In Kingdoms and Domains pp. 231–377. Academic Press, London.

Masta, S. E. and Boore, J. L. (2008). Parallel Evolution of Truncated Transfer RNA Genes in Arachnid Mitochondrial Genomes. Molecular Biology and Evolution, 25, 949–959. doi:10.1093/molbev/msn051.

Mikhailov, K. V., Slyusarev, G. S., Nikitin, M. A., Logacheva, M. D., Penin, A. A. and Aleoshin, V. V. (2016). The genome of *Intoshia linei* affirms orthonectids as highly simplified spiralians. Curr Biol, 26. doi:10.1016/j.cub.2016.05.007.

Nguyen, L.-T., Schmidt, H. A., von Haeseler, A. and Minh, B. Q. (2015). IQ-TREE: A Fast and Effective Stochastic Algorithm for Estimating Maximum-Likelihood Phylogenies. Molecular Biology and Evolution, 32, 268–274. doi:10.1093/molbev/msu300.

Ogino, K., Tsuneki, K. and Furuya, H. (2011). Distinction of cell types in *Dicyema japonicum* (phylum Dicyemida) by expression patterns of 16 genes. J Parasitol, 97. doi:10.1645/GE-2472.1.

Phillips, W. S., Brown, A. M. V., Howe, D. K., Peetz, A. B., Blok, V. C., Denver, D. R. and Zasada, I. A. (2016). The mitochondrial genome of Globodera ellingtonae is composed of two circles with segregated gene content and differential copy numbers. BMC Genomics, 17, 706. doi:10.1186/s12864-016-3047-x.

Podsiadlowski, L., Braband, A., Struck, T. H., von Döhren, J. and Bartolomaeus, T. (2009). Phylogeny and mitochondrial gene order variation in Lophotrochozoa in the light of new mitogenomic data from Nemertea. BMC Genomics, 10, 364. doi:1471-2164-10-364 [pii] 10.1186/1471-2164-10-364.

Robertson, H. E., Lapraz, F., Egger, B., Telford, M. J. and Schiffer, P. H. (2017). The mitochondrial genomes of the acoelomorph worms Paratomella rubra, Isodiametra pulchra and *Archaphanostoma ylvae*. Scientific Reports, 7, 1847. doi:10.1038/s41598-017-01608-4.

Schiffer, P., Robertson, H. and Telford, M. J. (2017). Molecular data from Orthonectid worms show they are highly degenerate members of phylum Annelida not phylum Mesozoa. bioRxiv.

Shao, R., Barker, S. C., Mitani, H., Aoki, Y. and Fukunaga, M. (2005). Evolution of duplicate control regions in the mitochondrial genomes of metazoa: a case study with Australian Ixodes ticks. Mol Biol Evol, 22. doi:10.1093/molbev/msi047.

Shao, R., Campbell, N. J. H. and Barker, S. C. (2001). Numerous Gene Rearrangements in the Mitochondrial Genome of the Wallaby Louse, Heterodoxus macropus (Phthiraptera). Molecular Biology and Evolution, 18, 858–865. doi:10.1093/oxfordjournals.molbev.a003867.

Shao, R., Kirkness, E. F. and Barker, S. C. (2009). The single mitochondrial chromosome typical of animals has evolved into 18 minichromosomes in the human body louse, *Pediculus humanus*. Genome Research, 19, 904–912. doi:10.1101/gr.083188.108.

Slyusarev, G. S. and Starunov, V. V. (2016). The structure of the muscular and nervous systems of the female *Intoshia linei* (Orthonectida). Organisms Diversity & Evolution, 16, 65–71. doi:10.1007/s13127-015-0246-2.

Telford, M. J., Herniou, E. A., Russell, R. B. and Littlewood, D. T. (2000). Changes in mitochondrial genetic codes as phylogenetic characters: two examples from the flatworms. Proceedings of the National Academy of Sciences, 97, 11359–11364. doi:10.1073/pnas.97.21.11359.

Tobias, Z. J. C., Yadav, A. K., Schmidt-Rhaesa, A. and Poulin, R. (2017). Intra- and interspecific genetic diversity of New Zealand hairworms (Nematomorpha). Parasitology, 144, 1026–1040. doi:10.1017/S0031182017000233.

Watanabe, K. I., Bessho, Y., Kawasaki, M. and Hori, H. (1999). Mitochondrial genes are found on minicircle DNA molecules in the mesozoan animal Dicyema. J Mol Biol, 286, 645–650. doi:10.1006/jmbi.1998.2523.

Webster, B. L. and Littlewood, D. T. J. (2012). Mitochondrial gene order change in Schistosoma (Platyhelminthes: Digenea: Schistosomatidae). International Journal for Parasitology, 42, 313–321. doi:https://doi.org/10.1016/j.ijpara.2012.02.001.

Weigert, A., Golombek, A., Gerth, M., Schwarz, F., Struck, T. H. and Bleidorn, C. (2016). Evolution of mitochondrial gene order in Annelida. Molecular Phylogenetics and Evolution, 94, 196–206. doi:https://doi.org/10.1016/j.ympev.2015.08.008.

